# Analyzing the Naming Conventions of Life Science Data Resources to Inform Human and Computational Findability

**DOI:** 10.1101/2025.10.02.680112

**Authors:** Heidi J. Imker, Hua Ou

**Affiliations:** University Library, University of Illinois Urbana-Champaign, Urbana, Illinois, USA; Epidemiology, Statistics, and Population Sciences Section, National Institute on Deafness and Other Communication Disorders (NIDCD), Bethesda, Maryland, USA

## Abstract

This study aimed to evaluate the names of life science data resources and consider the impacts on findability, a core feature of the FAIR (Findability, Accessibility, Interoperability, and Reusability) Principles. Utilizing a previously published list of unique data resources, we identified and validated data resources with both common and full names available (n = 1153). From this set, we analyzed characteristics of resource names to identify if any naming conventions have emerged organically. Additionally, since common names are often used in the absence of a resource’s full name, we performed a test to evaluate our ability to infer any meaning from common names. Our results highlight suboptimal naming practices and a wide-spread opaqueness in common names, which poses challenges to resource identification and retrieval by both human-and computationally-centric methods. These results are informative for those who establish and promote data resources as well as for those who search for data to use in individual research projects, develop data discovery systems, analyze the scientific literature, or assess research infrastructure. The findings underscore the value of findability in the FAIR Principles and the current efforts to develop infrastructure that supports more efficient communication and global connectedness.

## 1. Introduction

Work that began in the mid-20th century to aggregate and organize biological sequences in physical formats, such as Maragret Dayhoff’s *Atlas of Protein Sequence and Structure*, proved the immense value of developing such collections and paved the way for even greater impact through the coming computer revolution [1]. Today, centralized collections of digital data enable large-scale analyses that would not be possible without access to data collected across time, geographic boundaries, and individual studies. The far -reaching impacts of genomics could not have happened if, for example, DNA sequences were still only available as figures in individual articles. The transformative power of these collections has inspired the creation of thousands of new online data resources in the life sciences, also referred to as biodata resources. A major challenge for life science researchers today is simply finding which resources exist, knowing the data they contain, and how they may be used. This has intensified as funding agencies began implementing mandatory data sharing requirements, and researchers began seeking suitable repositories for data deposition.

The need to find data was codified in the FAIR Principles, with “F” standing for “findable”, as *“The first step in (re)using data is to find them. Metadata and data should be easy to find for both humans and computers. Machine-readable metadata are essential for automatic discovery of datasets and services, so this is an essential component of the FAIRification process*.” [2]. In addition to individual datasets, the discoverability of the data resources themselves—as the infrastructure that collects, organizes, and disseminates data collections—is also crucial [3]. Data resources provide valuable context and provenance, an essential component of assessing the appropriateness of data for any given task [4], [5]. Such resources are also sought out by those who may not intend to use the data directly, including those aiming to connect distinct resources (e.g., for portals, federated systems, etc.), assess the research landscape (e.g. to evaluate programs, policies, etc.), or provide research support (e.g., via data services). To enable findability across multiple audiences, the FAIR Principles emphasize the value of persistent identifiers (PIDs), and organizations that issue identifiers, such as DataCite, ensure that PIDs are associated with well-defined metadata in formal schema. In the ideal future state, PIDs with metadata that describe the key attributes of both the data resource itself and the data within will enable more facile discovery and reuse. This is a crucial step towards the AI-readiness that modern computational applications require [6], [7].

To achieve this future state, data resource creators must be aware of PIDs or organizations that issue PIDs must be aware of a specific resource. This awareness loop is difficult to close, but in either case, metadata fields require concrete values that may not always be codified or fixed. For example, one complex and often confusing value is also one of the most fundamental and critical—an entity’s title or name [8]. All data resources have a name, and these names are often so long that a shorter “common” name is also adopted. Common names exhibit a range of linguistic structures, formally referred to as extragrammatical morphologies, such as blends and acronyms, to represent portions of a longer “full” name. To complicate things further, both full and common names may have variations that arise purposely (e.g., on versioning of a resource) or spontaneously (e.g., by accident or whim). In a metadata schema, multiple names can be handled through fields dedicated to alternate names, but this still requires that these names be 1) clearly articulated and 2) documented in the metadata record. Between gaps in awareness and indefinite or incomplete attributes, it will be some time before all data resources have PIDs and comprehensive records.

This leaves those searching for data now, in an essentially “early-FAIR” era, with a problem—discovering data is difficult because data sources are distributed across the entire web and lack sufficient metadata for discoverability [5], [8], [9], [10]. Beyond PIDs, several efforts aim to enhance large-scale findability. Two examples incorporate Linked Data, a machine-readable, standard format for storing information that utilizes elements described by unique identifiers for interlinking. One example is the Data Catalog Vocabulary (DCAT), developed by the World Wide Web Consortium (W3C) to enhance representation of data catalogs on the web [11]. Another is Google’s Dataset Search, which was launched in 2018 and relies on use of schema.org elements on individual websites to identify data during web crawls [12]. These efforts are necessary since web crawling is resource-intensive, especially with the current scarcity of metadata on data resource websites. However, like PIDS, these global strategies require local awareness. Adoption of DCAT and schema.org is nascent, especially for data resources developed across the highly variable academic landscape. This has led to data-specific discovery specific strategies on the web. For example, Zhang and colleagues developed a method that began with user-supplied domain keywords to initiate a search followed by use of webpage classifiers, content analysis, and ranking to iteratively improve returns [10]. Additionally, even while enthusiasm for AI is exploding, current tools struggle to identify comprehensive lists of data resources because the models lack, among other things, the context and timeliness to serve as a “domain expert” [13]. Regardless, researchers searching for data now tend to follow familiar paths by entering keywords in general web browsers or similarly searching literature databases [4], [9], [12], [14]. That is, individuals’ strategies for data discovery are repurposed versions of their strategies for article discovery, despite the fact that data resources, let alone the data within, are not described and indexed to anywhere near the same degree. Given the challenges experienced by both human-and computationally-centric methods, we sought to learn current naming practices from existing data resources and if any norms, even informal, have emerged organically that can inform the findability of data as the ecosystem stands today.

This work is a follow-up to a study that used machine learning (ML) to identify several thousand biodata resources from the scientific literature [15]. In the present study, common and full names predicted by the ML model were validated and analyzed for length and common characteristics. Because of the prevalence of common names, we also performed an assessment of common name “clarity” to examine the extent to which these names can illuminate or obscure the purpose of resources. Our results provide insights into current naming practices and their impact on the findability of resources. They also lend themselves to suggestions that will enable better discoverability for the thousands of resources that currently exist as well as considerations for resources that have yet to be launched.

## 2. Results

### 2.1 Common name analysis

We analyzed the 1153 common names in our dataset and discovered that the average length was 5.36 characters, with a standard deviation of 2.06 (Fig 1). The shortest measured only two characters, while the longest common name spanned 40 characters. Among the 1153 common names, four-character names were the most prevalent (n = 321), followed by five characters (n = 262). Common names of three and six characters were nearly equal (n = 156 and n = 155, respectively). The 25th percentile contains names that are four characters or shorter, the median has names that are five characters, and the 75th percentile has names that are six characters or longer. Acronyms ranged from 3 to 9 characters in length for the vast majority (1100/1153) of resources, with a significant decrease in frequency for acronyms 10 characters or longer, making up < 5% of all resources.

**Fig 1.**
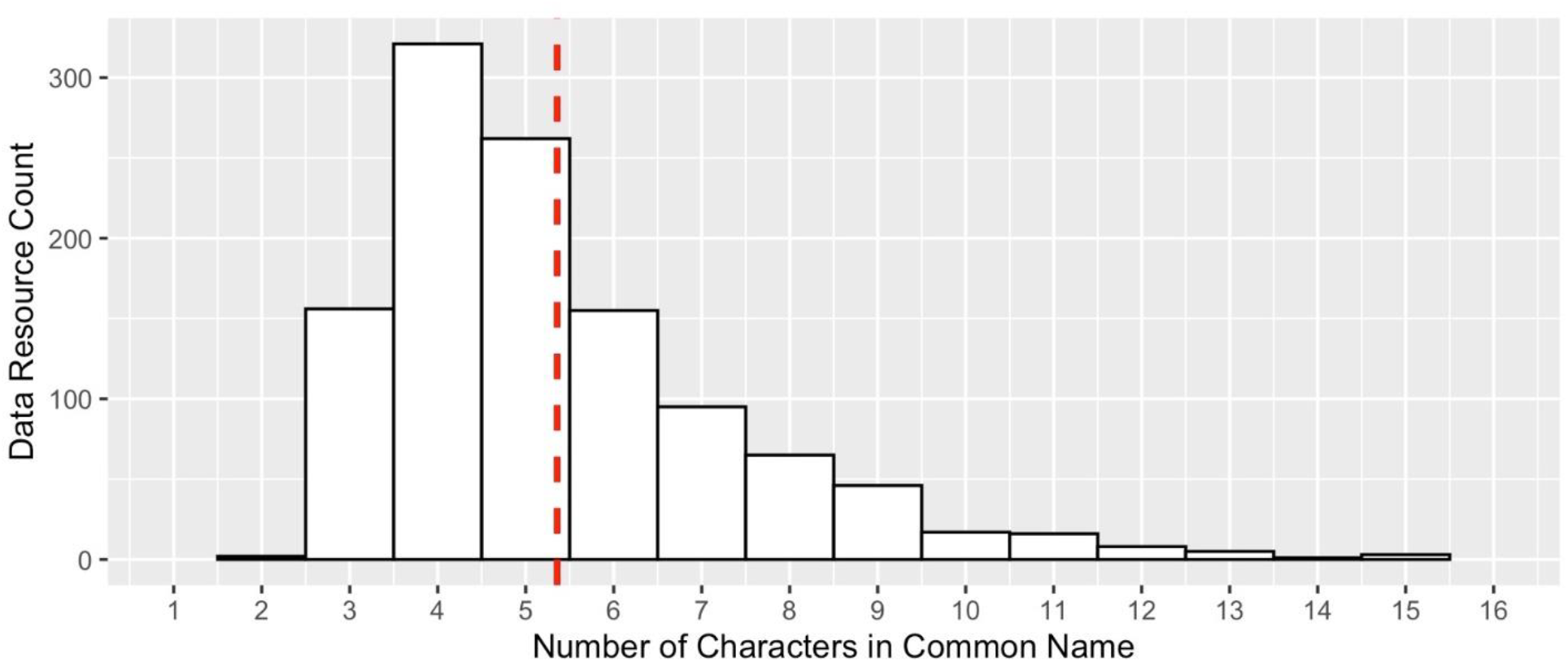
Common name length distribution. Histogram of 1153 common name lengths, in number of characters, with the mean value shown as a dashed red line.

In analyzing common prefixes and suffixes, we first explored a length of three characters as a typical affix length. We found 905 unique 3-character prefixes, with the most frequently found, “pla”, occurring 10 times. For suffixes, there were 697 unique 3-character values, with “mdb” the most frequent with 28 instances. However, the suffixes contained an obvious feature in that the last two characters were often “db”, including 18 of the top 20 most frequently occurring 3-character suffixes, prompting us to analyze just the last two characters as well. From this we found that “db” occurred in 261/1153 (23%) of the common names, 7 times more prevalent than the second most frequent 2-character e.orgnding, “gd”, which was found in 37 common names.

Although each data resource was unique within the list, we identified several instances of redundant common names. For example, MMDB was found three times, representing the distinct resources “Murine Microbiome Database,” “Molecular Modeling Database,” and “Magnaportheoryzae Microsatellite Database”. PED also occurred three times each, and 17 other acronyms, such as DGV, MPD, and PPD, were all found twice.

### 2.2 Agreement matrix for common name clarity

HO (the statistician) coded 1117/1153 (96.9%) common names as opaque, 35/1153 (3.03%) as translucent, and 1/1153 (0.087%) as transparent, while HJI (the biologist) coded 1012/1153 (87.8%) common names as opaque, 112/1153 (9.71%) as translucent, and 29/1153 (2.52%) as transparent.

Coders agreed on the vast majority of names (Fig 2), with both categorizing 1006/1153 (87.3%) as opaque, with 17/1153 (1.5%) agreed as translucent, and 0/1166 (0%) agreed as transparent.

**Fig 2.**
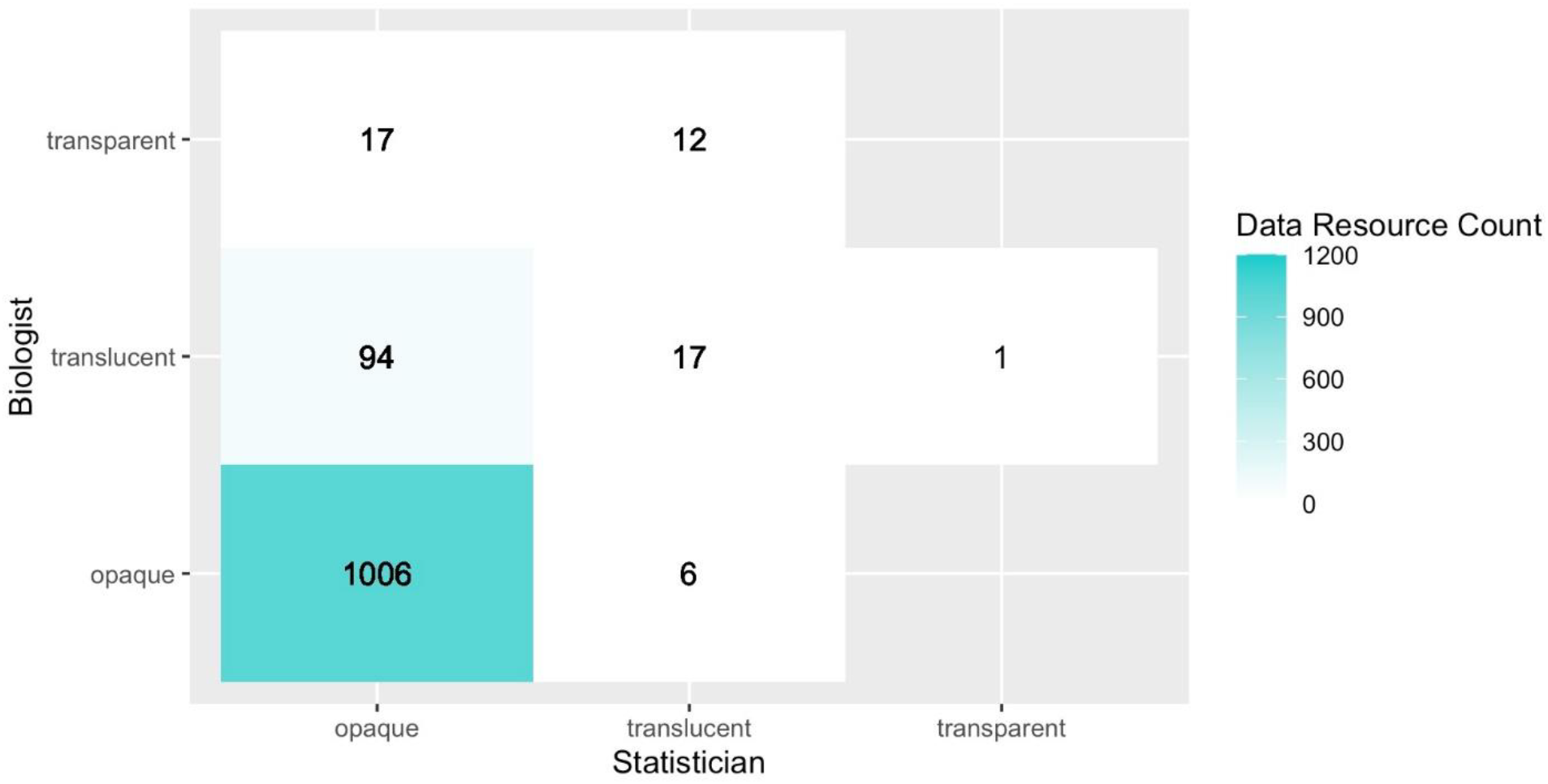
Agreement matrix for coded common names. The two authors from different professional backgrounds independently classified each common name as opaque, translucent, or transparent. The matrix shows the frequency of names in each pairwise coding combination, with perfect agreement along the diagonal.

### 2.3 Full name analysis

For the full set of 1153 corresponding full names, the average number of words per name was 4.49. The name lengths ranged from 1 to 14 words, with a standard deviation of 1.58 (Fig 3). The 25th percentile of names comprised 3 words or fewer, the median (50%) consisted of 4 words, and the 75th percentile included names that were 5 words or longer in length. The majority (1014/1153) of names fell between 2 and 6 words, with a pronounced decrease in frequency for names 7 words or longer, making up < 10% of all resources.

**Fig 3.**
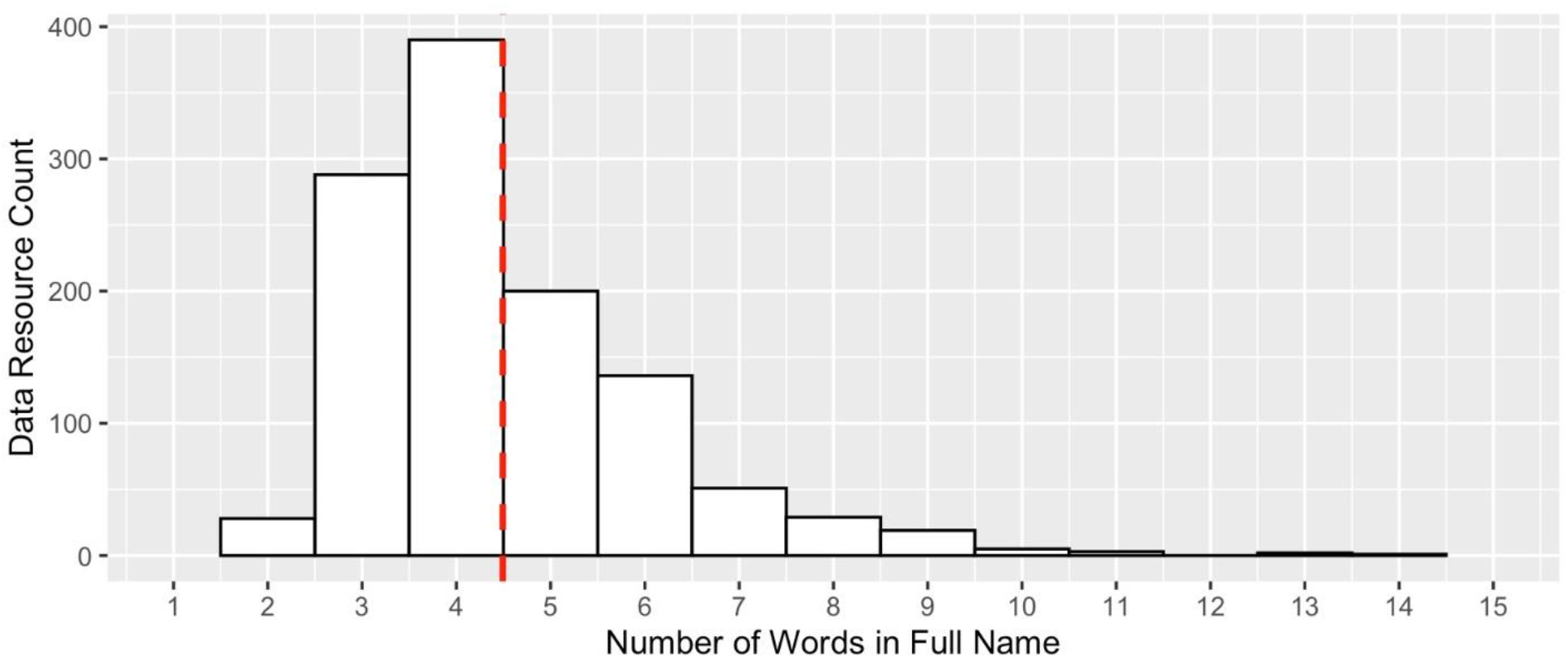
Full name length distribution. Histogram of 1153 full name lengths, in number of word, with the mean value shown as a dashed red line.

In analyzing the first and last words in full names, we found high variability in first words especially, which were in turn highly variable in the type of terms used. Examples include those with a starting word that indicated an organizational concept (e.g., “compendium”, “index”), an organism (e.g., “rat”, “silkworm”), a place name (e.g., “Leiden”, “China”), a location (e.g., “membrane”, “breast”), a disease (e.g., “cancer”, “Dengue”), a method (e.g., “crystallography”, “pharmacokinetics”), a molecular process (e.g., “diffusion”, “autophagy”), a molecular entity (e.g., “enzyme”, “carbohydrate”), and adjectives (e.g., “integrated”, “open”). In total, there were 684 unique first words, with the most frequently occurring being “database,” with 61 instances, followed by “human” with 41, and “plant” with 30. We found comparatively fewer unique last words with 304 in total. There were 559 instances of “database” as the last word, representing 48% of the 1153 resources, which was 13 times more common than the second most frequently found, “resource”, at 43 instances. In the top 10 most frequently occurring last words, only one, “protein” (10 instances) did not represent a type of organizational concept; the others were “atlas” (18), “network” (14), “base” (13), “portal” (13), “repository” (11), and “knowledgebase” (10).

## 3. Discussion

### 3.1 Data seeking audiences

Data are recognized as “boundary objects” that can bring multiple parties together [16], and community data repositories can be highly effective at bridging the “distance” between data creators and data reusers [17]. Traditionally, collaborative networks, conference experiences, and the scholarly literature have been key avenues for data discovery [4]. Undoubtedly, these avenues will remain crucial to developing data resource awareness, but they are not equally available to everyone [18]. Data resources, when discoverable, can further engage with a wider array of researchers by providing online access (often cost-free and unrestricted access) regardless of geographic, disciplinary, and organizational affiliation. Not only does this create a fairer landscape—an explicit goal of the Open Science movement [5], [19]—but larger user bases mean correspondingly greater reuse and potential to catalyze important discoveries. A wider range of use also demonstrate broader impacts and argue in favor of sustainable financial support.

Studies show that most researchers already consider themselves as belonging to more than one discipline and seek data from other disciplines [4], [12]. Beyond researchers’ existing tendency to cross boundaries, increased data sharing is tightly coupled with the concept of data reuse, and researchers are responding to competitions and new funding calls dedicated to supporting and championing reuse. Such efforts often encourage imaginative reuse, combining data, or use of lesser-known resources. For example, BioHackathons are annual events in Japan (since 2008) and Europe (since 2018), which aim to improve interoperability between data sources [20], [21]. In the US, the National Institutes of Health issued a Notice of Special Interest which included “Applications that use less or under-utilized data repositories or knowledgebases are encouraged.” [22]. Such opportunities provide another audience–those who have been incentivized to seek out unfamiliar data.

While direct users are understandably the focus of communication and engagement efforts, it’s important to note that audiences that may not reuse data themselves still seek data resources as they work to establish data discovery services, develop aggregation platforms, or assess the research ecosystem. Being discoverable to these groups is an even greater challenge since they may be even farther removed from a resource’s typical user community. Funders, publishers, and libraries have developed guides or lists of data repositories to assist researchers in their quest to find either data to reuse or appropriate places to deposit data (e.g., [23], [24], [25]). In the case of developing portals and knowledgebases, these efforts often aim to aggregate distinct data sources, and seeking out additional data provides a more comprehensive representation of a given topic. Funders also seek to know which data resources are created, used, or could be used by their fundees for the sake of policy implementation and program assessment, and others are interested in studying the research ecosystem’s data infrastructure as a whole (e.g., scientometric researchers). In all of these cases, it’s desirable to go beyond what is highly visible already, which leads to efforts to discover additional data resources.

### 3.2 Full names

There is a recognized need for alternative data search strategies, but the use of keywords and terms are still a core component of many discovery methods [14], [26]. In information retrieval research, searching for the specific, called a “look-up” or a “known item search”, relies on prior knowledge, and modern web search engines have been extraordinarily successful at quickly and easily meeting these well-defined needs, especially with the advent of technologies that incorporate semantic matches. It’s no wonder that researchers often start their data searches this way. However, discovery on the web via an “exploratory search” – when there is only a general idea of what is sought – is a continuous, investigative process that takes “strong” human interaction through iteration and interpretation [27]. This challenges the effectiveness of even the most sophisticated search tools. Beyond human-initiated searches, several computational methods also utilize keywords, including rule-based and dictionary NLP models. Additionally, terms may be included in the annotation guidelines used by human curators to create labeled data for training and fine-tuning in ML or AI applications [28], [29]. By evaluating data resource names, including the use of non-obvious terms, we hoped to inform both human-and computationally-centric strategies that build off of keywords.

The analysis of full names showed high variation in the first words, with most, but by no means all, being a scientific term. The last word is predominantly a term that represents, formally or informally, an organizational concept, with “database” the most frequently observed by far. This is unsurprising given the practical utility of databases as technical infrastructure and the corresponding prevalence of the term “database” within the life sciences. For example, *Nucleic Acids Research* has published a “Database Issue” since the early 1990s, which was so successful that a new journal, *Database: The Journal of Biological Databases and Curation*, was founded in 2009. The more illuminating result is the surfacing of alternative terms that can serve as a set of evidence-based synonyms, including “atlas”, “base”, “knowledgebase”, “network”, “portal”, “repository”, and “resource”, which all occurred 10 or more times. Additionally, relatively non-obvious and even less frequently used words, such as “compendium”, “directory”, “explorer”, “hub”, “index”, and “nexus” could further expand returns.

### 3.3 Common names

Abbreviations often serve as synonyms, or codes, for a more expanded word or phrase, and they are challenging to systematically study and categorize given their often informal, conversational origins and fuzzy, evolving use. In fact, clear definitions remain elusive in linguistics, but it is generally agreed, at the most basic level, that there are several types of abbreviations, including blends, consisting of word parts, and acronyms, consisting of characters. It is debated if an acronym must be pronounceable and form a word rather than be a simple initialism (e.g., radar vs EU); here we use the common, broader definition that either are acronyms [30], [31].

Acronyms are frequently used as common names for data resources. By utilizing letters or characters instead of word parts, they have the benefit of allowing the inventor to allude, however obliquely, to more terms using fewer characters and allowing for creativity, however subjectively. In writing for the *New Scientis*t in 1968, Eric Jamieson noted that university groups, and scientists in particular, are especially prone to creating acronyms [32]. In the following decades, Jamieson’s observation of what he termed “acronymania” has only intensified with recent measures showing that acronyms increased >3-fold in titles and >10-fold in abstracts between 1950 and 2019 [33]. Due to the high potential for confusion, there is no shortage of guidance on and criticisms of use (see [34]).

In our test to infer anything about a resource from its common name, the vast majority of common names we labeled as translucent or transparent were not strict acronyms, but various types of compounds and blends. Examples include AgeFactorDB, CottonGVD, DiseaseMeth, FlyRNAi, ShrimpGPAT, and Xenbase. These semantic clues directly related to the resource were informative. Notably, these abbreviations are distinct from acronyms that do form a word; here we found that these were often backronyms and semantically unrelated to a resource’s purpose. For example, COCONUT (COlleCtion of Open Natural prodUcTs), FROG-kb (Forensic Resource/Reference on Genetics-knowledge base), MAPS (Medicinal plant Activities), PICKLES (Pooled In vitro CRISPR Knockout Library Essentiality Screens), and SATuRN (Southern African Treatment Resistance Network). While such strategies may generate a memory aid, in the absence of context and consistent explanation, they may also be confusing and susceptible to redundancy. As early as 1997, SMART (which coincidentally was also found in the present study) was noted for redundancy [35], and a later study found 16 different clinical trials that used HEART as an acronym [36].

Computational detection may help identify and disambiguate resource names, but modern automated methods still depend on the information available. The study that generated our starting dataset identified over 3000 resources using transformer models to predict names during Named Entity Recognition (NER) tasks that used the titles and abstracts of articles classified as explicitly describing a given data resource. That study found that full names could be predicted for ∼40% of the resources, but these names carried lower F1 scores and were more likely to be fully or partially incorrect. This illuminates the inherent challenges for computational detection of context-rich full names, including their complexity relative to abbreviated names and the fact that they may not be present in the text at all, especially as a common name becomes the dominant form within a core community.

As a suffix, the abbreviation for database, “db”, was by far the most frequent feature observed among common names. Although ending in “db” was common enough at 23% that it may be helpful for recognizing a data resource as such, there is a trade-off. It exacerbates the potential for duplication since it effectively reduces variation, especially given the prevalence of short common names. In addition to the MMDB example above, another is HMDB, which is used for both the human metabolome database and the hydrocarbon monooxygenation database, despite 26^4^ (456,976) possibilities for 4 -character acronyms. While not frequent enough to serve as a meaningful clue, we note two common names inadvertently highlighted discovery issues while giving a nod to programmer humor by starting with “YA” for “Yet Another” (YADAMP: yet another database of antimicrobial peptides and YaTCM: Yet another Traditional Chinese Medicine database for drug discovery) [37]. Although we found little consistency otherwise among common names, these results are informative for setting up the parameters to recognize the presence of common names during web crawls or literature searches. Likewise, the character length range found can provide upper and lower limits during NER tasks.

### 3.4 Limitations

There are several limitations that impact the generalizability of this study. First, the inventory of biodata resources that provided the input for this study relied on mining the scientific literature via Europe PMC, which focuses on the life sciences. Thus, we do not know how well these results translate to data resources found in other domains or disseminated outside of the literature entirely. Second, that study included a targeted search for terms found within titles and abstracts, which could have resulted in over -representation of data resources with “data”, “database”, or “resource” within their names specifically, hence our emphasis on documenting alternative terms. Third, we focused exclusively on English language resources, and the use of other languages will undoubtedly impact data resource naming practices and findability. Fourth, classification of name clarity was the assessment of two individuals only, both of whom are proficient in English. Thus, we are not representative of all data seekers. Fifth, resource names are just one facet of data discovery data. For additional considerations, see [8], [9], [26].

## 4. Practical implications

Even as the FAIR movement matures for dataset-level metadata, where data came from will be obscured without clear, well-defined metadata at the collection level. This scenario played out for cultural heritage resources in the early 2000s, ultimately prompting the creation of collection level descriptions for Dublin Core, a widely used general purpose metadata schema [38], [39]. Similar collective-level description efforts for data resources are underway, e.g., BioDBcore for biological databases specifically [40] and, more generally, the Research Data Alliance’s new Recommendation for Common Descriptive Attributes of Research Data Repositories [8]. We hope that the work presented here shines another light on the importance of being able to discover data resources themselves, and one specific and critical metadata element—the resource’s name. The importance of systematic naming is well-understood by taxonomists and ontologists, who devote their careers to bringing clarity and cohesion to how entities are named, defined, and classified. Perhaps because the proliferation of data resources is a comparatively new phenomenon, naming practices have not been as systematic despite the role in discoverability.

For data resources, common names are often used for linguistic convenience, yet can slip into exclusive use and, in effect, take on the role of a brand name. However, compared to data resources, commercial brand names are less likely to be acronyms, or even abbreviations at all. This may be because they are formally understood and launched as brand names, taking into consideration, for example, conveying the purpose of the product [41]. In the commercial sector, strategic marketing experts recommend brand names that are memorable, meaningful, likable, transferable, adaptable, and protectable [42]. However, data resources often serve a highly complex and specific purpose, making it difficult to invent names that satisfy all of these qualities. While we acknowledge this challenge, inconsistencies in how data resource names are used hinders discoverability, which ultimately undermines data reuse across sectors. Given this, we encourage resource providers to consider the crucial role of naming and offer the following considerations explicitly related to resource names:

- For use of existing names
  - To provide more context for human and computational discovery, use full names frequently, e.g., within metadata for website markup and in the scientific literature
  - Guard against exclusive, overuse of common names, especially if it has no semantic relationship to the resource
- For determining future names
  - Common names will emerge, and the preemptive incorporation of branding best practices can inform their construction and make their use more effective
  - Set precedents at launch for consistently using resource name(s)

Establishing (and consistently using) clear and descriptive names will enable greater discovery immediately and as future “FAIRification” efforts create more efficient communication and global connectedness.

## 5. Methodology

### 5.1 Data source

The initial dataset was an inventory of 3112 unique biodata resources created using a fine -tuned version of BioMed-RoBERTa-RCT, a model previously pre-trained on the scientific literature for sequential sentence classification [43]. For the inventory, the fine-tuned model was able to predict a common name for 3078/3112 (98.9%) resources, a full name for 1243/3112 (39.9%), and both for 1166/3112 (35.9%) from the titles and abstracts of articles predicted to describe a biodata resource [15], [44]. Only resources with both names predicted were used in the present study so we could readily validate names of either type by cross-checking against each other. We examined each common-full name pair to assess the accuracy of the predicted names and further investigated by reviewing the associated articles and websites when needed. Inaccuracies and formatting errors were corrected where possible and invalid entries and near duplicates removed when not possible. This resulted in a final set of 1153 resources with validated common and full names. Both authors actively participated in verifying the accuracy and completeness of the names.

### 5.2 Analysis of common and full names

We applied a variety of analytical techniques to analyze and group data by examining common and full names using R and the packages tidyverse [45]. The purpose of this was to identify any inherent structures or similarities among either name type. Both common and full names were converted to lowercase before analysis and trimmed to remove leading or trailing whitespaces. Since most of the common names were acronyms, the analysis of common names included assessing how frequently each unique acronym appeared and an analysis of character lengths and affixes. For the analysis of full names, we calculated the number of words and analyzed the first and last words to explore patterns.

### 5.3 Clarity of common names

Given the relatively greater success of common name retrieval versus full name retrieval through machine learning methods, we assessed if common names contained information useful to humans about the nature of a given resource. The authors served as the coders and their varied backgrounds (HO -statistician; HJI -biologist) provided two distinct vantage points. The inherent subjectivity of this task reflects the real-world experience of data seekers. The 1153 commons names were independently reviewed by each author and coded as “transparent”, “translucent” or “opaque” where “transparent” indicated the coder had a clear sense the data resource’s topic, “translucent” indicated the coder could infer a topic, and “opaque” indicated that the coder could not infer anything about the resource. The coders’ assignments were analyzed in an agreement matrix to assess clarity overall.

## Funding

HJI wishes to thank the generous donor support as the Allen and Elaine Avner Professor in Interdisciplinary Research. The authors received no funding for this work.

## Data Availability Statement

The data, scripts, and associated documentation for this article are available on GitHub at https://github.com/1heidi/names and may be updated since the time of this manuscript’s preparation.

## Acknowledgements

The authors would like to thank MJ Han, Ken Schackart, and Chuck E. Cook for feedback on early drafts of this manuscript.

## Notes

https://github.com/1heidi/names

### Competing Interest Statement

The authors have declared no competing interest.

